# A unified method for rare variant analysis of gene-environment interactions

**DOI:** 10.1101/570226

**Authors:** Elise Lim, Han Chen, Josée Dupuis, Ching-Ti Liu

## Abstract

Advanced technology in whole-genome sequencing has offered the opportunity to comprehensively investigate the genetic contribution, particularly rare variants, to complex traits. Many rare variants analysis methods have been developed to jointly model the marginal effect but methods to detect gene-environment (GE) interactions are underdeveloped. Identifying the modification effects of environmental factors on genetic risk poses a considerable challenge. To tackle this challenge, we develop a unified method to detect GE interactions of a set of rare variants using generalized linear mixed effect model. The proposed method can accommodate both binary and continuous traits in related or unrelated samples. Under this model, genetic main effects, sample relatedness and GE interactions are modeled as random effects. We adopt a kernel-based method to leverage the joint information across rare variants and implement variance component score tests to reduce the computational burden. Our simulation study shows that the proposed method maintains correct type I error rates and high power under various scenarios, such as differing the direction of main genotype and GE interaction effects and the proportion of causal variants in the model for both continuous and binary traits. We illustrate our method to test gene-based interaction with smoking on body mass index or overweight status in the Framingham Heart Study and replicate the *CHRNB4* gene association reported in previous large consortium meta-analysis of single nucleotide polymorphism (SNP)-smoking interaction. Our proposed set-based GE test is computationally efficient and is applicable to both binary and continuous phenotypes, while appropriately accounting for familial or cryptic relatedness.

## 1. Introduction

Although genome-wide association studies (GWAS) have been successful in identifying genetic variants with strong association with disease, the variants evaluated in GWAS have been mostly restricted to common variants, typically defined as those with minor allele frequency (MAF) greater than 1 or 5%. Additionally, these identified variants only explain a small portion of disease heritability, which could be due to limited sample size and power in GWAS, and thus calls for performing meta-analysis.^1^ However, even in large scale GWAS meta-analyses, much of the heritability remains unexplained. For example, in GWAS meta-analysis of adult height in > 93,000 East Asians, they identified 98 loci at genome-wide significance that only explain about 9% of height heritability.^2^

One plausible explanation for missing heritability is the presence of rare variants, which are not analyzed in GWAS due to their low MAF. It is well known that rare variants are responsible for many Mendelian disorders, but they have not been fully implicated in complex diseases.^3,4^ With rapid advances and decreasing cost of whole-genome sequencing, attention has been shifted to investigating the potential role of low frequency variants in complex human disease etiology. There is now an abundance of growing empirical evidence that rare variants may be responsible for complex diseases, which might account for missing heritability.^4,5,6^ For example, Igartua et al.^7^ identified two novel rare variants associated with low-density lipoprotein cholesterol and high-density lipoprotein cholesterol with larger effects than the previously discovered variants within the known blood lipid associated loci. In another study by He et al., multiple rare variants were found to be associated with lower systolic blood pressure. Successes from rare variant association studies highlight the importance of rare variants to complex disease susceptibility.

When single variant tests are applied to rare genetic variants, they are severely underpowered, and hence several statistical methods have been developed to increase power. These methods are usually region-based tests that study the joint effects of rare variants in a specified region and can be broadly categorized into burden tests and non-burden tests. For burden tests, the cumulative effects of variants in a region are summarized into a single variable which is then tested for association with the trait of interest. Variants can be collapsed by summing up the number of rare alleles in a region or by using dichotomous variable to indicate whether an individual has any rare alleles in that region. Other extensions of burden tests have been developed. ^8,9,10^ Burden tests are most powerful when variants in consideration are causal and the direction of the effect on risk of the alternate/minor allele is the same. When the alternate/minor alleles of causal variants have effects on the phenotype in different direction, and/or the proportion of causal variants is small, burden tests suffer from low power.^4,12,13^ To address this issue, non-burden tests have been proposed that focus on aggregating individual test statistics. Among non-burden tests, the sequence kernel association test (SKAT), which is a popular method where variant information is summarized using a kernel function, uses variance component score test to evaluate the significance.^14^ SKAT is powerful when there are variants in mixed directions or when the proportion of causal variant is small, but it is less powerful than the burden tests when the majority of variants in a region are causal with effects in the same direction.^14,15^

Another plausible explanation for missing heritability is due to gene-environment (GE) interactions. Complex diseases are multifactorial and involve both genetics and environmental factors. Therefore, studying just the main effects, either genetic or environmental, cannot provide full insights into the biological mechanisms. Studying GE interactions can provide better insights into the biological mechanisms of complex diseases, which help to identify subgroups that are at high risk and eventually lead to better diagnostics and personalized treatments.^16,17^ Typically, GE tests require larger sample sizes than a main effects test to achieve appropriate power, and rare variant analysis also needs bigger sample compared to common variant analysis to maintain power. ^17,18,19^ Tzeng et al.^19^ developed similarity-based regression method (SimReg) to test GE interaction effects of rare variants for continuous traits. SimReg allows for covariate adjustments, models both main and interaction effects, and is computationally efficient. Zhao et al.^17^ extended SimReg to allow for binary traits for common or rare variants. Lin et al.^20^ introduced rare variant by environment interaction method using a variance component test under the generalized linear model framework. Chen et al.^18^ proposed two GE interaction tests (rareGE) and a joint test of main effect and interaction effect for rare variants using variance component score tests. In rareGE, genetic variants can be included as fixed or random effects for the interaction test and rareGE works for both binary and continuous traits. The aforementioned methods are only applicable to unrelated individuals and cannot correctly account for familial correlation in their models. Recently, Mazo Lopera et al.^21^ developed SNP-set GE interaction method for family data but it was proposed in the context of common variants.

In this article, we introduce a unified approach for GE interaction tests for rare variants (famGE) to correctly incorporate sample relatedness. Our approach can accommodate both binary and continuous traits in family data, eliminating the need to restrict samples to just unrelated individuals. We adopted a kernel-based method to leverage the joint information across rare variants. We assume main genetic variants, GE interaction term, and family correlation to be random effects and implemented a variance component score test in the generalized linear mixed model (GLMM) framework to reduce the computational burden.

The paper is organized as follows. In section 2, we introduce our notation, the GE interaction model, and the test statistic. In section 3, we conduct extensive simulations under various settings to assess type 1 error rates and power of our approach, comparing it to rareGE and the burden test. We apply our method in testing gene by smoking interaction on body mass index (BMI), using family data from the Framingham Heart Study (FHS) in section 4. We discuss our findings of our approach in section 5.

## 2. Methods

*Notations and model settings for interaction test for family data*

We first introduce our notation and assumptions before we derive our test statistic. Assuming a sample of size *n*, let *Y_i_* be the phenotype (binary or continuous) for individual *i* with and *E*(*Y_i_*) = *μ_i_*, and *var*(*Y_i_* = *ϕv*(μ_i_), where *ϕ* is the dispersion parameter (1 for binary and Poisson data), and *v*(·) is the variance function. Let ***X**_i_* = (1,*E*_*i*_,*X*_*i*2_, …,*X*_*i*2_(*p*-1))^T^ be a p x 1 vector of covariates including the intercept and the environmental variable *E_i_*, and ***G**_i_* = (*G*_*i*1_, *G*_*i*2_, …, *G_iq_*)^*T*^ be a q x 1 vector of genetic main effects, and *d_i_* be the random effect of familial correlation. We consider the following GLMM for testing GE interactions:

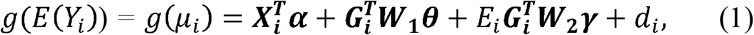

where *g*(·) is the link function, ***α*** is a p x 1 vector associated with the fixed covariate ***θ*** effects, is a q x 1 vector of random effects for the genetic variants, ***γ*** is a q x 1 vector of random effects for GE interaction, and ***W*_1_** and ***W*_2_** are q x q diagonal matrices with weights for genetic main effects and GE interaction effects, respectively. The matrices ***W*_1_** and ***W*_2_** can be any positive semi-definite matrices. We can rewrite this model in matrix notation as:

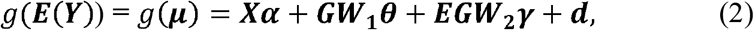

where *μ* = [*μ*_1_ … *μ_n_*]^T^ is the mean vector, ***X*** = [***X*_1_** … ***X**_n_*]^T^ is an n x p covariate matrix, ***G*** = [***G*_1_** … ***G_n_***]^*T*^ is an n x q genotype matrix, ***EG*** = [***EG*_1_** … ***EG_n_***]^*T*^ is an n x q GE interaction matrix, and ***d*** = [*d*_1_ … *d_n_*]^*T*^ is an n x 1 vector for the random effects of familial correlation. For the three random effects (genetic variants, GE interactions, and relatedness in families respectively), we assume that

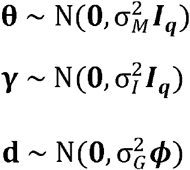

where ***ϕ*** is twice the n x n kinship matrix from family relationships obtained from a pedigree or an empirical kinship calculated from genotype data to account for cryptic relatedness. The random effects ***θ, γ***, and ***d*** are assumed to be independent from each other. If ***γ*** is treated as a fixed effect, we would perform a *q* degrees of freedom score test, but this test can lead to power loss when *q* is large.^12^ By assuming that ***γ*** follows a normal distribution with mean 0 and variance 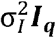, the null hypothesis for the interaction test: *H_0_:* ***γ*** = 0 is equivalent to testing 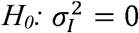 with a variance component score test.^12,14^ Score tests only require fitting the model under the null hypothesis, so they are more computationally efficient.

### Estimation and hypothesis testing

To fit the null model for binary traits, we use the penalized quasi-likelihood method. Because the integrated quasi-likelihood function for binary traits contains a high dimensional integral that is intractable, we use the Laplacian method to approximate the integral. To estimate the parameters, we define the linear working vector under the null hypothesis ***Y*_0_ = *Xα* + *GW*_1_ θ + *d* + Δ(*y −μ*)**, where **Δ** = *diag*{*g^′^*(*μ_i_*), and let 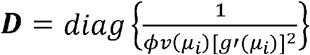. We iteratively fit the working vector to estimate the parameters until convergence to obtain our restricted maximum likelihood (or maximum likelihood) estimates.^22^

Let 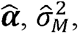, and 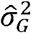 be the restricted maximum likelihood (or maximum likelihood) estimates under the null hypothesis. We can then calculate 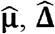, and 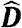 and 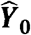 using the aforementioned restricted maximum likelihood (or maximum likelihood) estimates. The restricted maximum quasi-likelihood function is defined as:

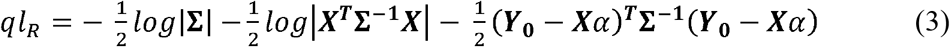

where 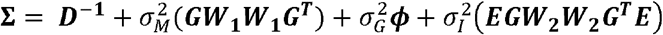.

To derive the score test for 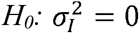, we take the first derivative of the quasi-likelihood with respect to 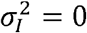,

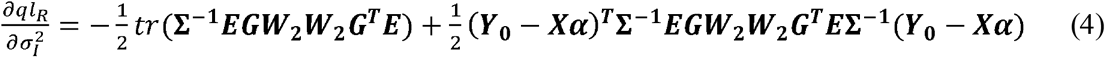

The first term in equation 4 is fixed and independent of the phenotype. We follow the same rationale in the derivation of the SKAT score statistic and take twice the second term to be our test statistic^14,23^

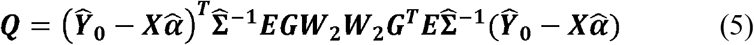

where 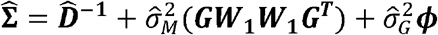.

Under the null hypothesis, 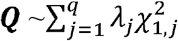, where *λ_j_*’s are eigenvalues of the matrix 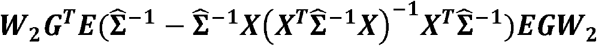.^14,24^ See Appendix 1 for a detailed derivation.

## 3. Simulation studies

### Simulation settings for type I error evaluation

We performed simulation studies to evaluate type I error rates for our proposed approach. For the null simulation study, we first considered the scenario where there are main genotype effects but no GE interaction effects. To simulate the genotypes, we used SeqSIMLA software and used reference sequence based on 1000 Genomes Project for European populations.^25^ We used 7 pedigrees from FHS with family membership ranging from 120 to 640 (2030 individuals in total). In the simulated genotypes, we chose a region that spans from 1,100 base pairs to 1,140 base pairs in chromosome 1. To simulate our phenotype, we varied the proportion of low frequency (MAF<5%) causal SNPs included in our model to 20% to 40% to 60% and 80% for each of 20,000 replicates. We considered both continuous and binary phenotypes. For each of 20,000 replicates, we simulated phenotype datasets from

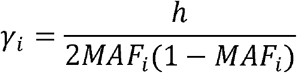

where ***age*** was generated from a normal distribution with mean of 50 and standard deviation equal to 5, ***sex*** was generated from a Bernoulli distribution with probability 0.5, ***smoke*** was generated from a Bernoulli distribution with probability 0.5, and ***ϵ*** was generated from a standard normal distribution. Family correlation, *d*, is from a multivariate normal distribution with means 0 and covariance 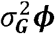, where 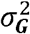 is set to 1 ***ϕ*** and is twice the kinship matrix. ***γ*** consist of effect sizes for the causal SNPs. For binary traits, we simulated the continuous traits and set the lower 80% as controls (0’s) and the upper 20% as cases (1’s). We simulated variants where the directions of the effect on risk of the minor allele are both same and opposite and the effect sizes were determined by

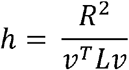

where MAF is the minor allele frequency of SNP *i*. The constant *h* is calculated as
where *R*^2^, the proportion of variance explained by the causal SNPs, was fixed at 1% for causal SNPs with effect sizes in same direction and 5% for causal SNPs with effect sizes in opposite directions. The correlations between the SNPs are in matrix *L*, and *v* is a vector that indicates the direction of the SNP effects. We performed three tests for the type I error comparison: our proposed method that correctly accounts for familial correlation (famGE), rareGE where the main genetic effects are treated as random and family correlation is ignored (rareGE), and the burden test using GLMMs to account for familial correlation (BT). The summary variable used in the burden test was created using an indicator of whether or not the rare allele was present in the testing region. We simulated 20,000 replicates and used Wu weights, which are the beta density function with parameters 1 and 25 evaluated at the MAF of the variants, for famGE and rareGE tests.^14^

### Simulation settings for power analysis

To assess power, we simulated data under the alternative hypothesis, where we include a gene by smoking interaction effect in addition to the main genotype effects. Similar to the type I error simulation, genotypes were simulated using the SeqSIMLA software and we varied the direction of genetic main effect and the proportion of causal SNPs with MAF less than 5%. We simulated 10,000 phenotype datasets from

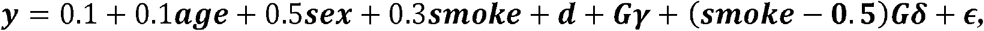

where age, sex, and smoke, family correlation, and error terms were generated from the same distribution described in the type I error simulation study and the genotype effects were determined the same way as in our null simulation study. Interaction effects ***δ*** were generated from a normal distribution with mean 2 and standard deviation of 0.3. We considered the scenarios where directions of the main effects ***δ*** and the interaction effects on the risk of the minor allele are same and opposite. Negative interaction effects were simulated the same way as above except we multiplied the effects by -1. To test binary outcomes, we set the lower 80% of the simulated continuous outcome to be the controls and the upper 20% to be the cases. For power comparison, we also performed a burden test, where the summary variable for each individual was created using an indicator of whether or not the rare allele was present in the testing region.

### Results for type 1 error simulations

**Table 1** includes the type 1 error results for famGE, rareGE, and BT at significance levels, α, of 0.05, 0.01, and 0.001 from 20,000 simulation replicates. Both famGE and BT have correct type 1 error rates at all three α levels for both continuous and binary traits. When familial correlation is not appropriately taken into account in the model, rareGE test suffers from type 1 error inflation, which is more pronounced in binary traits. FamGE has valid type 1 error rates under various scenarios, such as differing the direction of main genotype effects and increasing the proportion of causal variants in the model for both continuous and binary traits. Because rareGE does not have correct type 1 error rate when familial correlation is not taken into account, we did not include rareGE in the power comparison in the next section.

**Table 1.** Comparison of type 1 errors based on 20,000 replications.

| Scenarios (+/-/0) | $\alpha$ level | Continuous | | | Binary | | |
| --- | --- | --- | --- | --- | --- | --- | --- |
|  |  | famGE<sup>a</sup> | rareGE<sup>b</sup> | BT<sup>c</sup> | famGE<sup>a</sup> | rareGE<sup>b</sup> | BT<sup>c</sup> |
| 4/0/16 | 0.05 | 0.0496 | 0.0530 | 0.0501 | 0.0503 | 0.0551 | 0.0507 |
|  | 0.01 | 0.0100 | 0.0110 | 0.0102 | 0.0102 | 0.0125 | 0.0105 |
|  | 0.001 | 0.0010 | 0.0170 | 0.0010 | 0.0012 | 0.0018 | 0.0010 |
| 8/0/12 | 0.05 | 0.0488 | 0.0521 | 0.0499 | 0.0484 | 0.0557 | 0.0492 |
|  | 0.01 | 0.0100 | 0.0103 | 0.0100 | 0.0097 | 0.0134 | 0.0095 |
|  | 0.001 | 0.0010 | 0.0011 | 0.0010 | 0.0011 | 0.0019 | 0.0009 |
| 12/0/8 | 0.05 | 0.0495 | 0.0520 | 0.0499 | 0.0482 | 0.0554 | 0.0498 |
|  | 0.01 | 0.0094 | 0.0120 | 0.0096 | 0.0106 | 0.0122 | 0.0090 |
|  | 0.001 | 0.0010 | 0.0015 | 0.0010 | 0.0008 | 0.0015 | 0.0010 |
| 16/0/4 | 0.05 | 0.0504 | 0.0538 | 0.0500 | 0.0484 | 0.0570 | 0.0500 |
|  | 0.01 | 0.0091 | 0.0116 | 0.0099 | 0.0097 | 0.0141 | 0.0094 |
|  | 0.001 | 0.0010 | 0.0016 | 0.0010 | 0.0010 | 0.0020 | 0.0010 |
| 2/2/16 | 0.05 | 0.0480 | 0.0525 | 0.0500 | 0.0497 | 0.0556 | 0.0506 |
|  | 0.01 | 0.0095 | 0.0120 | 0.0101 | 0.0094 | 0.0116 | 0.0097 |
|  | 0.001 | 0.0010 | 0.0014 | 0.0011 | 0.0011 | 0.0016 | 0.0011 |
| 4/4/12 | 0.05 | 0.0499 | 0.0533 | 0.0502 | 0.0484 | 0.0552 | 0.0489 |
|  | 0.01 | 0.0093 | 0.0105 | 0.0100 | 0.0097 | 0.0116 | 0.0096 |
|  | 0.001 | 0.0010 | 0.0019 | 0.0010 | 0.0010 | 0.0014 | 0.0009 |
| 6/6/8 | 0.05 | 0.0493 | 0.0541 | 0.0494 | 0.0488 | 0.0561 | 0.0488 |
|  | 0.01 | 0.0103 | 0.0110 | 0.0104 | 0.0092 | 0.0137 | 0.0100 |
|  | 0.001 | 0.0010 | 0.0014 | 0.0010 | 0.0011 | 0.0017 | 0.0011 |
| 8/8/4 | 0.05 | 0.0502 | 0.0531 | 0.0493 | 0.0489 | 0.0555 | 0.0479 |
|  | 0.01 | 0.0090 | 0.0111 | 0.0099 | 0.0097 | 0.0127 | 0.0094 |
|  | 0.001 | 0.0010 | 0.0017 | 0.0012 | 0.0010 | 0.0016 | 0.0010 |

+/-/0: number of variants with main genotype effect sizes that are positive, negative, and neutral
famGE<sup>a</sup>: our proposed method accounting for family correlation
rareGE<sup>b</sup>: interaction test ignoring family correlation
BT<sup>c</sup>: burden test

### Results for power simulations

**Figure 1** shows the power comparison for famGE and BT for continuous traits at *α* = 0.001 from 10,000 replicates. When the proportion of causal variants is low, BT has lower power than famGE (**1A**). This is expected because burden tests are powerful when a large proportion of causal variants are included in the region. When the proportion of causal variants increases, we see that powers for both BT and famGE increase but we see a larger power increase in BT compared to famGE. However, when variants have interaction effects in different direction, we observe a power drop, with power close to 0 for BT, whereas famGE shows increasing power as proportion of causal variants increases (**1B**). FamGE maintains fairly consistent power across different scenarios, regardless of the direction of the main effects.

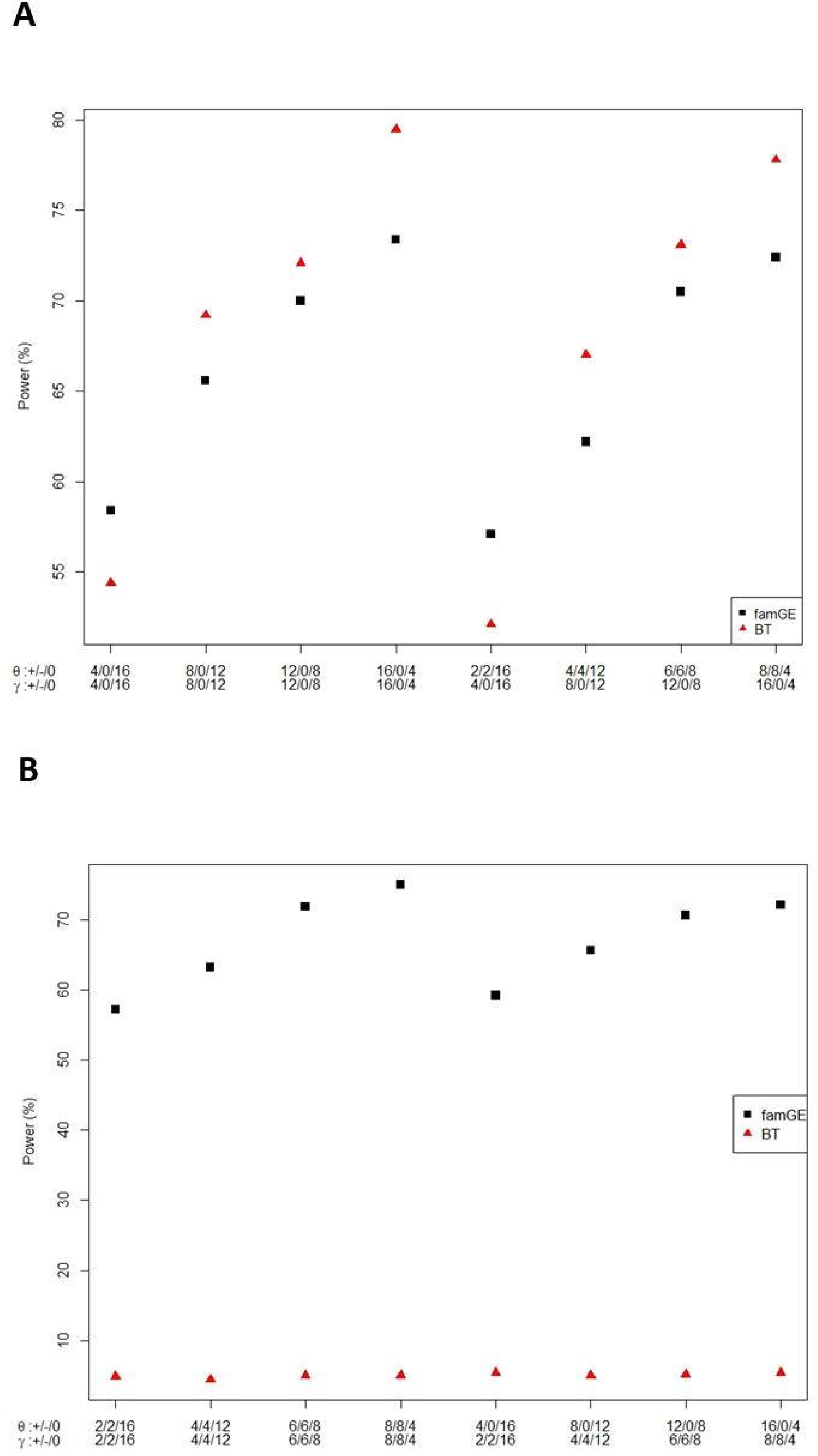

**Figure 2** shows power comparisons of famGE and BT for binary traits at *α* = 0.001 from 10,000 replicates. We come to the same conclusion as the results seen from continuous traits. Even though burden tests have higher power than famGE in the case where interaction effects are in the same direction, with the exception where there are 4 causal variants and 16 neutral variants in the model, famGE is able to maintain high power regardless of the direction or the proportion of variants (**2A**). The burden test significantly loses power in the presence of both trait-increasing and decreasing alleles, whereas performance of famGE is robust across different scenarios (**2B**).

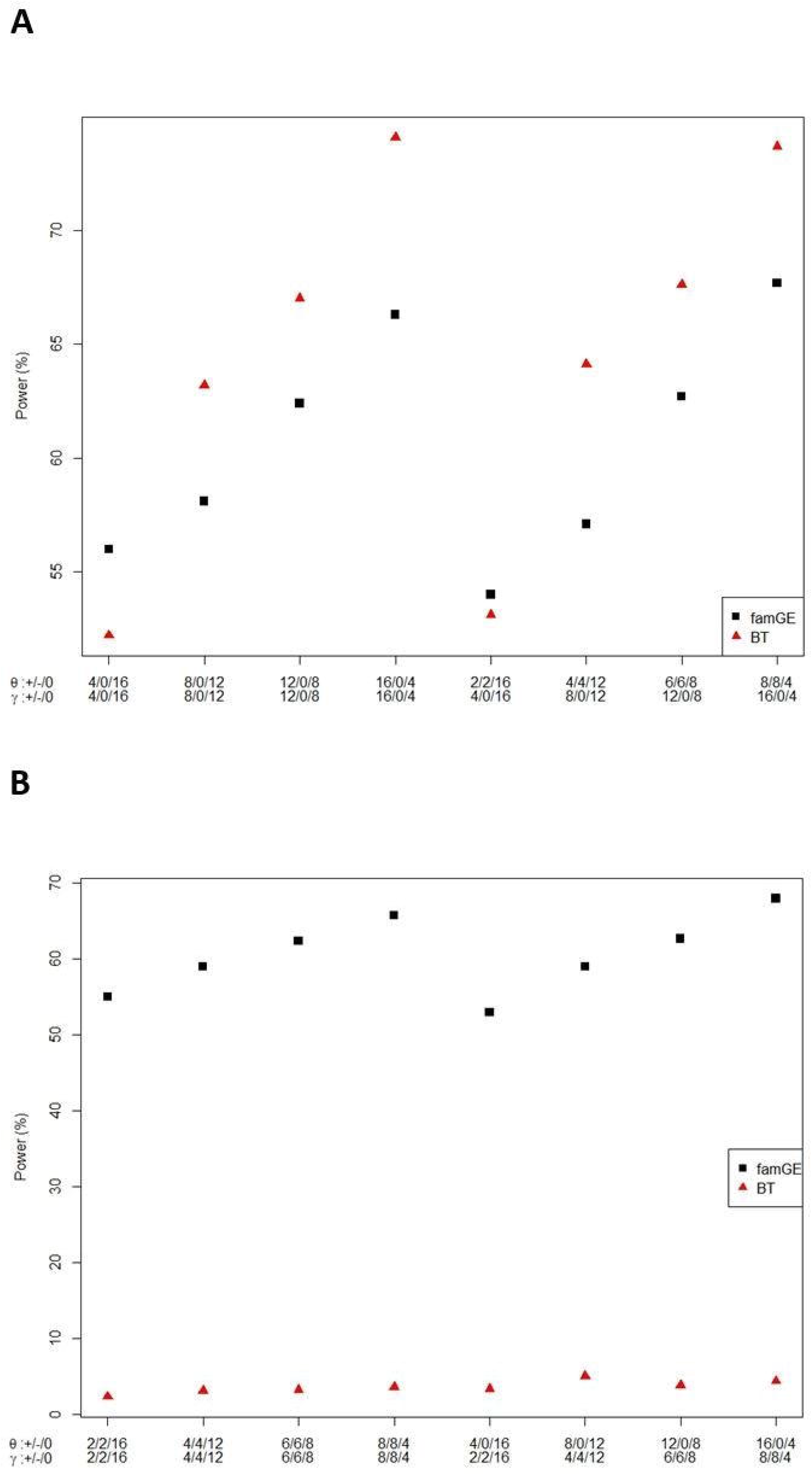

## 4 Application to the Framingham Heart Study

In the real data application, we illustrate our method to test gene-based interaction with smoking on a quantitative trait, BMI, and a dichotomous trait, overweight status (BMI ≥ 25), using participants from the Framingham Heart Study (FHS). FHS is a longitudinal cardiovascular cohort study that was initiated in 1948 in Framingham, Massachusetts. The study began with 5209 participants (Original Cohort) in 1948 and in 1971, 5124 children of the original cohort and their spouses (Offspring Cohort) were enrolled. A total of 4095 additional participants, who were the grandchildren of the original cohort, were recruited in the third generation cohort (Gen 3 Cohort) in 2002.^26^ To reflect ethnic diversity in Framingham, 917 additional non-white participants were recruited to form the Omni Cohort.^27^

Obesity is a world-wide problem that can lead to serious health problems, such as high blood pressure, type 2 diabetes, heart disease. Conventionally, BMI has been used to measure obesity. GWAS have identified numerous loci associated with BMI, but whether these genetic effects are modulated by environmental factors has not been extensively investigated.^28,29^ Recently, Justice et al.^30^ investigated the effect of smoking on genetic susceptibility to obesity in a large consortium meta-analysis of 241,258 individuals and reported two common variants reaching genome-wide significance, rs12902602 and rs336396 near the *CHRNB4* and *INPP4B* genes respectively. Here we evaluated whether there are modification effects of smoking status on genetic risk from these two loci for obesity with rare or less frequency variants.

We analyze genotype data from the Illumina V1.0 Exome Chip and select variants with MAF less than 5%. We adjust for age, sex, cohort, first two principal components, and smoking main effect in our model. Our data consists of 596 individuals from the Original Cohort (Exam 21), 2547 from the Offspring Cohort (Exam 8), 3868 from the Gen 3 Cohort (Exam 1), and 177 from the Omni Cohort (Exam 1). There are 3264 males (45.4% males), 6063 non-smokers (84.4% non-smokers), and their age ranges from 19 to 85 (median = 49). We test for gene x smoking interaction by treating BMI as continuous trait or dichotomizing BMI at 25 into a binary trait, which classified 4502 individuals as overweight and 2686 individuals in the normal range. A total of 7188 individuals are included in the gene x smoking interaction analysis.

We consider two genes, Cholinergic Receptor Nicotinic Beta 4 (*CHRNB4*) and Inositol Polyphosphate-4-Phosphatase (*INPP4B*). **Table 2** summarizes the analysis results for the two aforementioned genes. We include variants that are annotated as either stop-gain/loss, splice, or missense. Using famGE, we find *CHRNB4* to be statistically significant at *α* = 0.025 for both continuous (p-value = 0.0063) and dichotomized (p-value = 0.0063) BMI, but no gene x smoking interaction were identified for gene *INPP4B*.

**Table 2.**
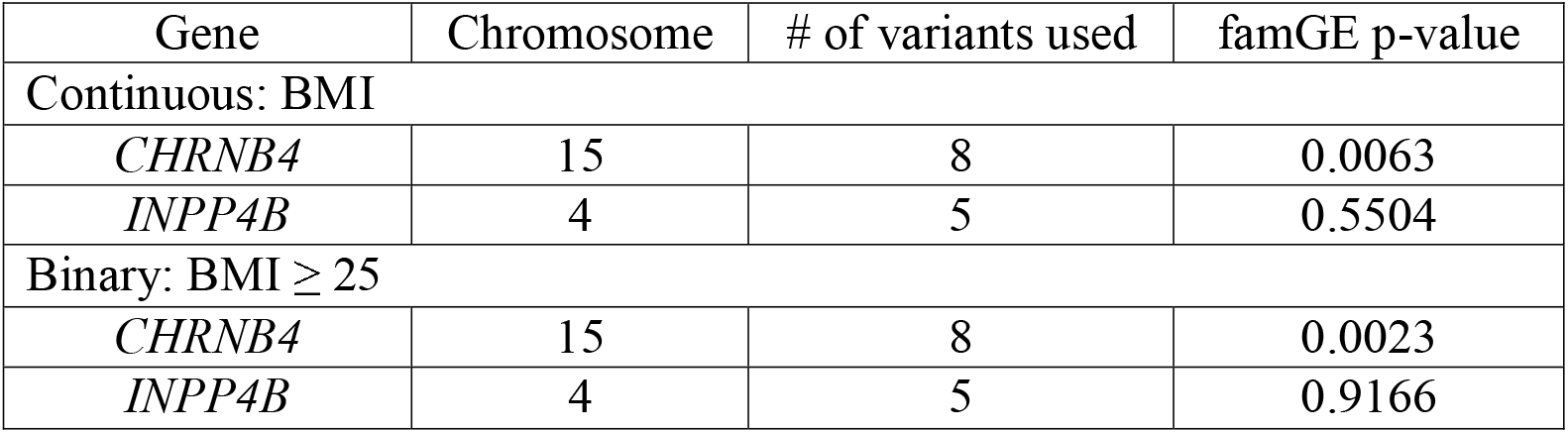
Association results of gene by smoking interaction on continuous trait: BMI and binary trait: overweight status (BMI ≥ 25) for two previously identified loci using the Framingham Heart Study.

## 5. Discussion

In this article, we develop a unified method called famGE to detect GE interactions of a set of rare variants under the GLMM framework. This proposed approach can accommodate both binary and continuous traits in family data or samples with cryptic relatedness. Additionally, famGE allows weighting variants differently based on prior information such as allele frequency. Under this model, we treat the genetic variants, familial correlation, and GE interaction effects as random effects and implement a variance component score test, which only requires fitting the null model, and thus, reduces computational burden. Our simulation studies show that famGE maintains correct type 1 error and high power under various scenarios. An R package famGE is available upon request.

Ignoring familial correlation when using linear or logistic regression to analyze family data lead to inflated type I error.^23^ One way to resolve this issue is to select a subset of unrelated individuals, but this may substantially reduce the sample size and lead to power loss. In the case of rare variants, especially for GE interaction test, it is important to retain as many samples as possible to have appropriate power. A preferable method is to use GLMMs that account for familial correlation as a random effect and thus, eliminates the need to restrict to unrelated individuals. In famGE, kinship coefficients can be obtained either from a pedigree or an empirical kinship matrix. Because the empirical kinship matrix can account for cryptic relatedness, it is advantageous to use the empirical kinship to estimate the level of relatedness among individuals.^14^

The proposed famGE method models the main genotypes as random effects in order to reduce the number of parameters that need to be estimated. If one wishes to model the main effect as fixed effects, the derivations of the new test statistic follows the same framework as the statistic for famGE with random genotype main effects. When the number of variants included in the model is large, however, there is a potential for multicollinearity when fitting the model. It is also shown that modeling the genetic main effects as fixed leads to slightly inflated type 1 error rate at less stringent α levels when the number of variants included in the testing region is large.^18^ Therefore, modelling genetic main effects as random effects is preferable.

From the two genes we test in the real data application, *CHRNB4* shows statistical significance in interacting with smoking on BMI. The *CHRNB4* gene has been reported to be associated with higher BMI in never smokers and lower BMI in current smokers, implying that genetics variants may influence BMI via the weight-reducing effects of smoking in opposite directions.^31^ *INPP4B* gene is a novel locus identified in the metaanalysis by Justice et al.^30^ but to our knowledge, this finding has not yet been replicated in other studies. We cannot exclude the possibility that there are no rare variants interacting with smoking in *INPP4B* gene.^30^ However, it is also possible that reduced power due to limited sample size in our study compared to that of in the meta-analysis restrict us from finding a significant association.

With advanced development and decreasing cost of sequencing technology, rare and low frequency variants have become more easily accessible, and there is growing evidence that they are implicated in complex diseases. Therefore, more attention has been brought to analyzing and developing rare variants methods. GE interaction may explain part of the missing heritability, provide insight into etiology of disease, identify subgroups in the population that are at high risk, and help develop personalized treatments. Our proposed approach provides a powerful, robust and efficient test to identify GE interaction of rare variants for both continuous and binary traits in the sample with related or independent participants.

## Acknowledgements

This work was in part supported by NIH grants: T32 GM 074905 (EL), R00 HL 130593 (HC) and U01 HL 120393 (HC), U01 DK078616 (JD and CTL), R01 HL120393 (CTL), R01 DK089256 (EL and CTL), and R01 HL118305 (EL and CTL).

## Appendix 1: Derivation of famGE

We used a generalized linear mixed model (GLMM) framework to derive the famGE statistic. We consider the following gene-environment (GE) model:

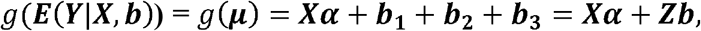

where ***b*_1_** = **GW_1_θ, *b*_2_** = **EGW_2_*γ***, and ***b*_3_** = **d** and ***Z*** = **[*I_n_, I_n_, I_n_*]**, 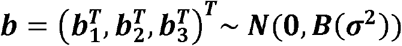, where 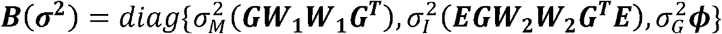 depends on an unknown vector of variance components ***σ*^2^**.

For subject *i*, the quasi-likelihood given random effects ***b*** is defined by:

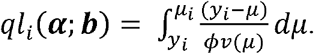

The integrated quasi-likelihood function used to estimate (***α, σ*^2^**) is written as

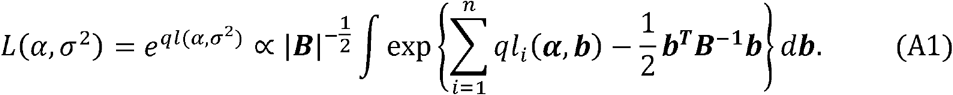

After applying the Laplacian transformation for integral approximation, the log of A1 becomes

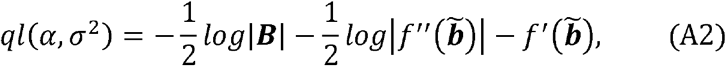

where 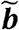 is the solution to

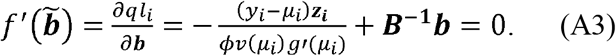

For canonical link functions, the second partial derivative with respect to b is equal to

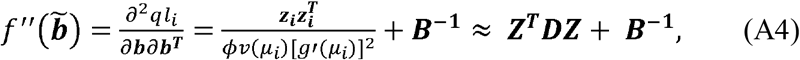

where 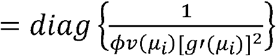.

Combining equations (A3) – (A4), equation (A2) becomes

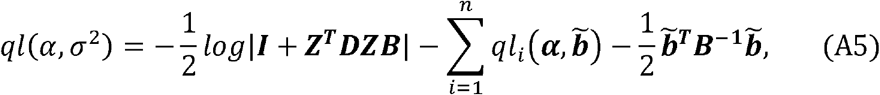

where 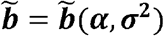 is the solution to

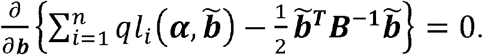

Defining **Δ** = diag{*g′*(*μ_i_*)} and assuming that the weight matrix ***D*** vary slowly as a function of the mean^22^, we maximize by differentiating with respect to **α** and **b**:

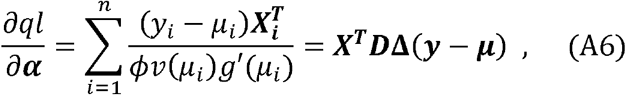

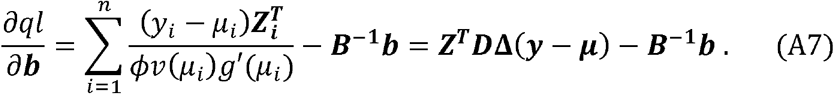

Defining the working vector ***Y*_0_** = ***Xα* + *Zb* + Δ(*y* – *μ*)**, solutions to A6 and A7 can be expressed as an iterative solution to the system

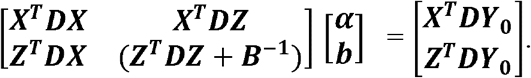

The test statistic for 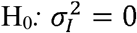 is

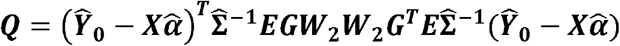

where 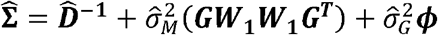

