## Supplementary 1 for "A unified method for rare variant analysis of gene-environment interactions"

**Supplementary 1: Derivation of famGE**

We used a generalized linear mixed model (GLMM) framework to derive the famGE statistic. We consider the following gene-environment (GE) model:

$g\boldsymbol{(E}\left( \boldsymbol{Y|X,b} \right)$**) *=*** $g\left( \boldsymbol{\mu} \right)\boldsymbol{=X\alpha+}\boldsymbol{b}_{\boldsymbol{1}}\boldsymbol{+}\boldsymbol{b}_{\boldsymbol{2}}\boldsymbol{+}\boldsymbol{b}_{\boldsymbol{3}}\boldsymbol{=X\alpha+Zb,}$

where $\boldsymbol{b}_{\boldsymbol{1}}\mathbf{=G}\mathbf{W}_{\mathbf{1}}\boldsymbol{\theta}$, $\boldsymbol{b}_{\boldsymbol{2}}\mathbf{=EG}\mathbf{W}_{\mathbf{2}}\boldsymbol{\gamma}$, and $\boldsymbol{b}_{\boldsymbol{3}}\mathbf{=d}$ and **Z = [**$\boldsymbol{I}_{\boldsymbol{n}}\boldsymbol{,}\boldsymbol{I}_{\boldsymbol{n}}\boldsymbol{,}\boldsymbol{I}_{\boldsymbol{n}}$**],** $\boldsymbol{b=}\left( \boldsymbol{b}_{\boldsymbol{1}}^{\boldsymbol{T}}\boldsymbol{,}\boldsymbol{b}_{\boldsymbol{2}}^{\boldsymbol{T}}\boldsymbol{,}\boldsymbol{b}_{\boldsymbol{3}}^{\boldsymbol{T}} \right)^{\boldsymbol{T}}\boldsymbol{\sim N(0,B}\left( \boldsymbol{\sigma}^{\boldsymbol{2}} \right)\boldsymbol{)}$**,** where $\boldsymbol{B}\left( \boldsymbol{\sigma}^{\boldsymbol{2}} \right)\boldsymbol{=}diag\{\sigma_{M}^{2}\left( \boldsymbol{G}\boldsymbol{W}_{\boldsymbol{1}}\boldsymbol{W}_{\boldsymbol{1}}\boldsymbol{G}^{\boldsymbol{T}} \right)\boldsymbol{,}\sigma_{I}^{2}\left( \boldsymbol{EG}\boldsymbol{W}_{\boldsymbol{2}}\boldsymbol{W}_{\boldsymbol{2}}\boldsymbol{G}^{\boldsymbol{T}}\boldsymbol{E} \right)\boldsymbol{,}\sigma_{G}^{2}\boldsymbol{\phi}\}$ depends on an unknown vector of variance components $\boldsymbol{\sigma}^{\boldsymbol{2}}$.

For subject *i,* the quasi-likelihood given random effects ***b*** is defined by:

${ql}_{i}\left( \boldsymbol{\alpha};\boldsymbol{b} \right)\boldsymbol{=}\int_{y_{i}}^{\mu_{i}} \frac{(y_{i}-\mu)}{\phi v(\mu)}d\mu$.

The integrated quasi-likelihood function used to estimate ($\boldsymbol{\alpha,}\boldsymbol{\sigma}^{\boldsymbol{2}}\boldsymbol{)}$ is written as

$$L\left( \alpha,\sigma^{2} \right)\boldsymbol{=}e^{ql(\alpha,\sigma^{2}\boldsymbol{)}}\boldsymbol{\propto}\left| \boldsymbol{B} \right|^{\boldsymbol{-}\frac{1}{2}}\int\exp\left\{ \sum_{i=1}^{n} ql_{i}\left( \boldsymbol{\alpha,b} \right)\boldsymbol{-}\frac{1}{2}\boldsymbol{b}^{\boldsymbol{T}}\boldsymbol{B}^{\boldsymbol{-1}}\boldsymbol{b} \right\}d\boldsymbol{b.}(A1)$$

After applying the Laplacian transformation for integral approximation, the log of A1 becomes

$$ql\left( \alpha,\sigma^{2} \right)\boldsymbol{=-}\frac{1}{2}\log\left| \boldsymbol{B} \right|-\frac{1}{2}\log\left| f^{''}\left( \tilde{\boldsymbol{b}} \right) \right|-f^{'}\left( \tilde{\boldsymbol{b}} \right)\boldsymbol{,}(A2)$$

where $\tilde{\boldsymbol{b}}$ is the solution to

$f^{'}\left( \tilde{\boldsymbol{b}} \right)\boldsymbol{=}$ $\frac{\partial ql_{i}}{\partial\boldsymbol{b}}=$ $-\frac{\left( y_{i}-\mu_{i} \right)\boldsymbol{z}_{\boldsymbol{i}}}{\phi v\left( \mu_{i} \right)g'(\mu_{i})}+\boldsymbol{B}^{\boldsymbol{-1}}\boldsymbol{b}=0$. (A3)

For canonical link functions, the second partial derivative with respect to b is equal to

$f^{''}\left( \tilde{\boldsymbol{b}} \right)\boldsymbol{=}$ $\frac{\partial^{2}ql_{i}}{\partial\boldsymbol{b}\partial\boldsymbol{b}^{\boldsymbol{T}}}=$ $\frac{\boldsymbol{z}_{\boldsymbol{i}}\boldsymbol{z}_{\boldsymbol{i}}^{\boldsymbol{T}}}{\phi v\left( \mu_{i} \right){[g'(\mu_{i})]}^{2}}+\boldsymbol{B}^{\boldsymbol{-1}}\approx\boldsymbol{Z}^{\boldsymbol{T}}\boldsymbol{DZ}+ \boldsymbol{B}^{\boldsymbol{-1}}\boldsymbol{,}$	(A4)

where $=diag\left\{ \frac{1}{\phi v\left( \mu_{i} \right){[g'(\mu_{i})]}^{2}} \right\}$ .

Combining equations (A3) – (A4), equation (A2) becomes

$$ql\left( \alpha,\sigma^{2} \right)\boldsymbol{=-}\frac{1}{2}\log\left| \boldsymbol{I+}\boldsymbol{Z}^{\boldsymbol{T}}\boldsymbol{DZB} \right|-\sum_{i=1}^{n} ql_{i}\left( \boldsymbol{\alpha,}\tilde{\boldsymbol{b}} \right)\boldsymbol{-}\frac{1}{2}{\tilde{\boldsymbol{b}}}^{\boldsymbol{T}}\boldsymbol{B}^{\boldsymbol{-1}}\tilde{\boldsymbol{b}}\boldsymbol{,}(A5)$$

where $\tilde{\boldsymbol{b}}\boldsymbol{=}\tilde{\boldsymbol{b}}\boldsymbol{(\alpha,}\boldsymbol{\sigma}^{\boldsymbol{2}}\boldsymbol{)}$ is the solution to

$\frac{\partial}{\partial\boldsymbol{b}}\left\{ \sum_{i=1}^{n} ql_{i}\left( \boldsymbol{\alpha,}\tilde{\boldsymbol{b}} \right)\boldsymbol{-}\frac{1}{2}{\tilde{\boldsymbol{b}}}^{\boldsymbol{T}}\boldsymbol{B}^{\boldsymbol{-1}}\tilde{\boldsymbol{b}} \right\}=0$.

Defining $\boldsymbol{\Delta}=diag\left\{ g'(\mu_{i}) \right\}$ and assuming that the weight matrix ***D*** vary slowly as a function of the mean ^22^, we maximize by differentiating with respect to $\boldsymbol{\alpha}\mathrm{and}\boldsymbol{b}$:

$$\frac{\partial ql}{\partial\boldsymbol{\alpha}}=\sum_{i=1}^{n} \frac{\left( y_{i}-\mu_{i} \right)\boldsymbol{X}_{\boldsymbol{i}}^{\boldsymbol{T}}}{\phi v\left( \mu_{i} \right)g'(\mu_{i})}=\boldsymbol{X}^{\boldsymbol{T}}\boldsymbol{D}\boldsymbol{\Delta}\left( \boldsymbol{y-\mu} \right) , (A6)$$

$$\frac{\partial ql}{\partial\boldsymbol{b}}=\sum_{i=1}^{n} \frac{\left( y_{i}-\mu_{i} \right)\boldsymbol{Z}_{\boldsymbol{i}}^{\boldsymbol{T}}}{\phi v\left( \mu_{i} \right)g'(\mu_{i})}-\boldsymbol{B}^{\boldsymbol{-1}}\boldsymbol{b}=\boldsymbol{Z}^{\boldsymbol{T}}\boldsymbol{D}\boldsymbol{\Delta}\left( \boldsymbol{y-\mu} \right)-\boldsymbol{B}^{\boldsymbol{-1}}\boldsymbol{b .}(A7)$$

Defining the working vector $\boldsymbol{Y}_{\boldsymbol{0}}=\boldsymbol{X\alpha+Zb+}\boldsymbol{\Delta}\left( \boldsymbol{y-\mu} \right)$, solutions to A6 and A7 can be expressed as an iterative solution to the system

$\left[ \begin{matrix} \boldsymbol{X}^{\boldsymbol{T}}\boldsymbol{DX} & \boldsymbol{X}^{\boldsymbol{T}}\boldsymbol{D}\boldsymbol{Z} \\ \boldsymbol{Z}^{\boldsymbol{T}}\boldsymbol{DX} & \boldsymbol{(Z}^{\boldsymbol{T}}\boldsymbol{D}\boldsymbol{Z+}\boldsymbol{B}^{\boldsymbol{-1}}\boldsymbol{)} \end{matrix} \right]$ $\left[ \begin{matrix} \boldsymbol{\alpha} \\ \boldsymbol{b} \end{matrix} \right]$ = $\left[ \begin{matrix} \boldsymbol{X}^{\boldsymbol{T}}\boldsymbol{D}\boldsymbol{Y}_{\boldsymbol{0}} \\ \boldsymbol{Z}^{\boldsymbol{T}}\boldsymbol{D}\boldsymbol{Y}_{\boldsymbol{0}} \end{matrix} \right]$.

The test statistic for H_0_*:* $\sigma_{I}^{2}=0$ is

$${\boldsymbol{Q=}\left( {\hat{\boldsymbol{Y}}}_{\boldsymbol{0}}-\boldsymbol{X}\hat{\boldsymbol{\alpha}} \right)}^{\boldsymbol{T}}{\hat{\boldsymbol{\Sigma}}}^{\mathbf{-1}}\boldsymbol{EG}\boldsymbol{W}_{\boldsymbol{2}}\boldsymbol{W}_{\boldsymbol{2}}\boldsymbol{G}^{\boldsymbol{T}}\boldsymbol{E}{\hat{\boldsymbol{\Sigma}}}^{\mathbf{-1}}\boldsymbol{(}{\hat{\boldsymbol{Y}}}_{\boldsymbol{0}}-\boldsymbol{X}\hat{\boldsymbol{\alpha}})$$

where $\hat{\boldsymbol{\Sigma}}\boldsymbol{=}{\hat{\boldsymbol{D}}}^{\boldsymbol{-1}}\boldsymbol{+}\hat{\sigma}_{M}^{2}\left( \boldsymbol{G}\boldsymbol{W}_{\boldsymbol{1}}\boldsymbol{W}_{\boldsymbol{1}}\boldsymbol{G}^{\boldsymbol{T}} \right)\boldsymbol{+}\hat{\sigma}_{G}^{2}\boldsymbol{\phi}$**.**
